# Synaptic RTP801 Contributes to Motor Learning Dysfunction in Huntington’s Disease

**DOI:** 10.1101/2020.04.07.030080

**Authors:** Núria Martín-Flores, Leticia Pérez-Sisqués, Jordi Creus-Muncunill, Mercè Masana, Sílvia Ginés, Jordi Alberch, Esther Pérez-Navarro, Cristina Malagelada

**Affiliations:** Department of Biomedicine, Faculty of Medicine, University of Barcelona, Barcelona, 08036 Catalonia, Spain; Institut de Neurociències, University of Barcelona, Barcelona 08036, Catalonia, Spain; IDIBAPS-Institut d’Investigacions Biomèdiques August Pi i Sunyer, Barcelona, 08036 Catalonia, Spain; Centro de Investigación Biomédica en Red sobre Enfermedades Neurodegenerativas (CIBERNED), Barcelona, 08036 Spain

**Keywords:** Huntington’s disease, plasticity, motor learning, mTOR, RTP801, Akt

## Abstract

RTP801/REDD1 is a stress responsive protein that mediates mutant huntingtin (mhtt) toxicity in cellular models and is up regulated in Huntington’s disease (HD) patients’ putamen. Here, we investigated whether RTP801 is involved in motor impairment in HD by affecting striatal synaptic plasticity.

Ectopic mhtt was over expressed in cultured rat primary neurons. The protein levels of RTP801 were assessed in homogenates and crude synaptic fractions from human postmortem HD brains and mouse models of HD. Striatal RTP801 expression was knocked down with adeno-associated viral particles containing a shRNA in the R6/1 mouse model of HD and motor learning was then tested.

Ectopic mhtt elevated RTP801 in synapses of cultured neurons. RTP801 was also up regulated in striatal synapses from HD patients and mouse models. Knocking down RTP801 in the R6/1 mouse striatum prevented motor learning impairment. RTP801 silencing normalized the Ser473 Akt hyperphosphorylation by downregulating Rictor and it induced synaptic elevation of calcium permeable GluA1 subunit and TrkB receptor levels, suggesting an enhancement in synaptic plasticity.

These results indicate that mhtt-induced RTP801 mediates motor dysfunction in a HD murine model, revealing a potential role in the human disease. These findings open a new therapeutic framework focused on the RTP801/Akt/mTOR axis.

## INTRODUCTION

Huntington’s disease (HD) is an autosomal-dominant neurodegenerative disorder caused by a CAG expansion in the exon 1 of the *Huntingtin* gene. This expansion encodes for a mutant form of the huntingtin (htt) protein that has been traditionally pointed out as responsible for the specific loss of medium-sized spiny neurons in the human striatum^1–3^. However, the expanded CAG RNA was identified also as toxic and an active contributor to the HD pathogenesis^4^. HD manifests a triad of signs including severe motor dysfunction with involuntary movements (chorea), cognitive impairment and neuropsychiatric symptoms. Even though mutant htt severely affects striatal neurons, other areas, such as cortex, hippocampus, amygdala or cerebellum, display synaptic alterations, atrophy and/or neuronal death^5,6^.

Although neuronal death does not occur until late stages of HD, abnormal synaptic plasticity and neuronal dysfunction are the main early pathogenic events that lead to neurodegeneration^7–9^. Due to the early aberrant synaptic function, observed both in the human and the mouse pathology, HD is considered a synaptopathy^10–12^. In this regard, one of the pathways that controls synaptic plasticity is the mechanistic target of rapamycin (mTOR) pathway, since it regulates translation and more notably, local protein synthesis at the spines^13,14^. Importantly, mTOR pathway is also involved in cytoskeleton remodeling to ensure proper formation and function of dendritic spines^15^.

mTOR kinase is the central component of mTOR complex (mTORC) 1 and 2. Both complexes share protein partners, but they have unique elements that define substrates’ specificity and therefore, functionality. First, mTORC1 binds specifically to Raptor and controls mostly protein synthesis and autophagy. Second, mTORC2 specifically binds to Rictor and phosphorylates the Serine 473 residue (Ser473) in Akt kinase to mediate neuronal survival. Among other functions, mTORC2 controls actin polymerization and, as a consequence, it is required for LTP and LTD induction to mediate synaptic strength^16,17^.

Synaptic plasticity is impaired in HD patients^18^ and mouse models^19–21^. These plastic alterations correlate well with mTOR/Akt signaling axis impairment. For example, several HD mouse models show increased phosphorylation levels of striatal mTOR and the mTORC2 substrate Akt at the Ser473 residue^22–24^. Interestingly, PHLPP1 (PH domain leucine-rich repeat protein phosphatase 1), the phosphatase that dephosphorylates the Ser473 residue in Akt^25,26^, is decreased in the putamen of HD patients and in the striatum of HD mice^23^. Moreover, mTORC2-regulator protein Rictor, but not mTORC1-regulator protein Raptor, is increased in the striatum of HD mouse models and in the putamen of HD patients^24^. This evidence suggested the activation of a compensatory mechanism to counteract mhtt toxicity promoting cell survival. However, the exacerbation on the phosphorylation status of mTOR pathway components could eventually be counterproductive for synaptic plasticity^27,28^.

In our previous work, we described RTP801/REDD1, an mTOR/Akt modulator, as a mediator of mhtt toxicity in *in vitro* models of HD. We also showed that RTP801 was elevated in the putamen and iPSC from HD patients^29^. RTP801, by interacting with TSC1/2 complex, regulates Rheb promoting its GDP-bound form. Rheb-GDP is not able to promote mTOR kinase activity of both mTORC1 and 2 complexes, inactivating S6 kinase and Akt activities, as readouts of mTORC1 and mTORC2, respectively. This mechanism was described to mediate neuronal death in Parkinson’s disease (PD) models^30,31^. However, since in HD there is an hyperphosphorylation of mTOR in the striatum of both HD mouse models and patients, the mechanism by which RTP801 contributes to the specific mTOR/Akt axis over activation and how this is translated to impaired plasticity has not been elucidated yet. For this reason, here we investigated whether mhtt-induced RTP801 increase could alter mTOR/Akt signaling at a synaptic level and mediate motor impairment in HD.

## RESULTS

### Ectopic mhtt increases RTP801 at soma and dendritic spines in rat cortical primary neurons

We have previously shown that overexpression of the pathogenic exon-1 of mhtt elevates RTP801 protein levels in rat cortical neuronal cultures^29^. Here, we extended these studies by analyzing whether RTP801 increase takes place in the dendrites. The use of cortical primary cultures facilitated this analysis since striatal mono-cultures generally produce morphologically immature MSNs with low densities of dendritic spines^32–35^.

Hence, rat cortical primary neurons (DIV13) were transfected with eGFP, Q25, Q72- or Q103-htt-expressing plasmids. Twenty-four hours after transfection, neurons were stained against endogenous RTP801 and dendritic F-actin puncta were visualized with phalloidin (Figure 1A). As indicated (yellow arrowheads), RTP801 was found within the dendritic spines. The overexpression of exon-1-mhtt (Q72 and Q103 pathogenic forms) induced a reduction in spine density (Figure 1B) along with an increase in somatic RTP801 (Figure 1C), as previously reported^29,36^. Importantly, mhtt overexpression increased around 50% the levels of RTP801 protein at the remaining dendritic spines (Figure 1D).

**Figure 1.**
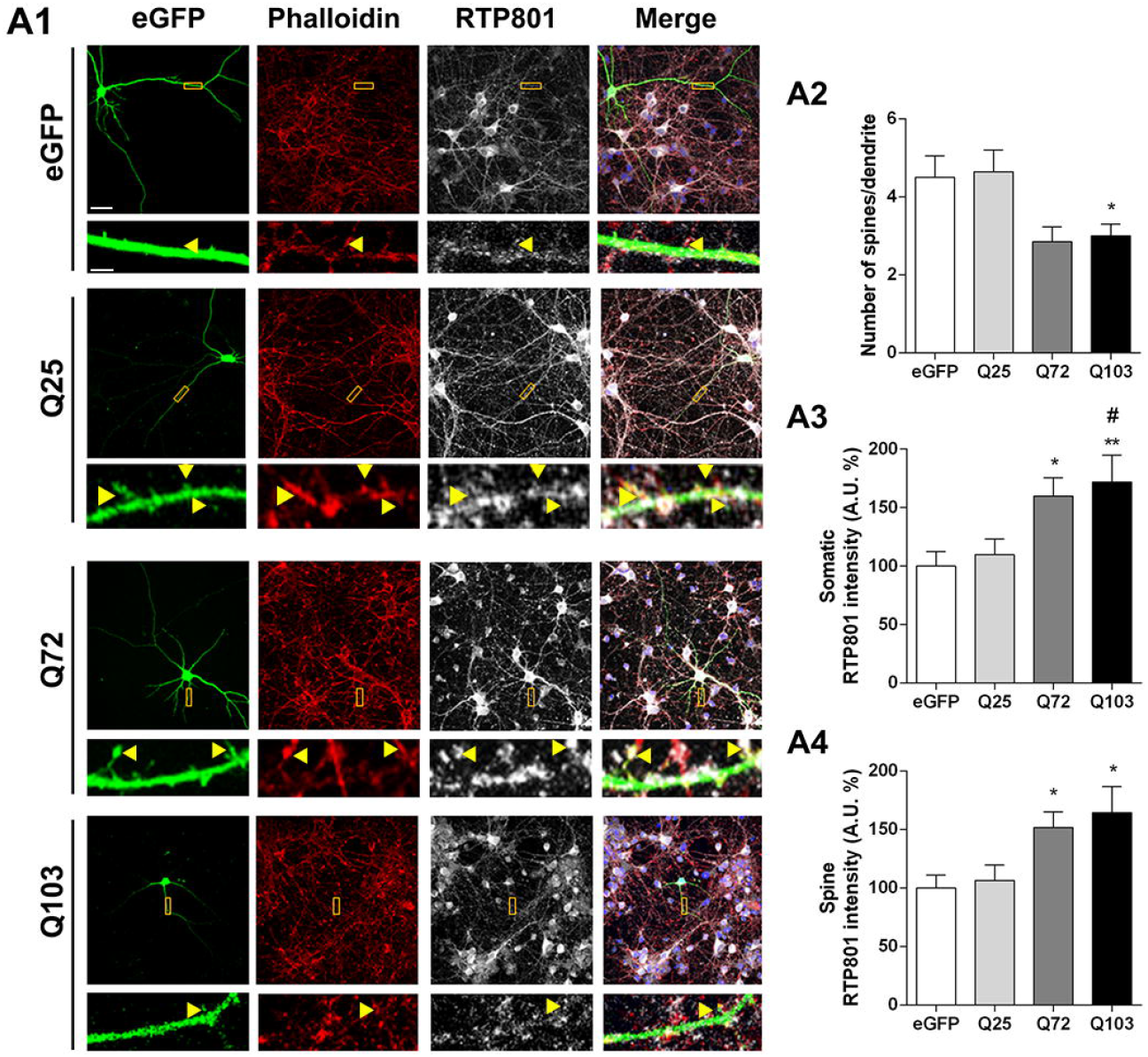
Overexpression of exon-1 mhtt increases RTP801 protein levels in dendrites and in synaptic spines. **(A1)** Rat cortical cultures at DIV13 were transfected with eGFP, Q25, Q72 or Q103 plasmids. Twenty-four hours after transfection cultures were fixed and stained against GFP (in green), RTP801 (in grey) and phalloidin (in red) to visualize the actin cytoskeleton. Images were acquired by confocal microscopy. Yellow rectangles show digital zoom of dendrites with spines and yellow arrows show RTP801 staining at the puncta. Scale bar, 10μm and in higher magnification images, 1.5μm. **(B)** Graphs display the number of spines scored for each 15 μm dendrite-length and RTP801 staining intensity quantification at **(C)** the neuronal soma and at **(D)** the spines. Data are shown as percentage of RTP801 intensity (mean ± SEM) of four independent experiments and were analysed with One-way ANOVA followed by Dunnett’s multiple comparisons test (\**P*□ 0.05 and \*\**P*□0.01 *vs*. eGFP and #*P*□0.05 *vs*. Q25, n=13 dendrites for eGFP, n=10 dendrites for Q25, Q72 and Q103 per experiment).

### RTP801 is increased in the synaptic compartment in the putamen of HD patients

Our previous work showed that RTP801 was elevated in whole putamen lysates from HD patients. To investigate further whether the increase in RTP801 also extends to synaptic contacts, we isolated crude synaptosomes from the putamen of non-affected (CT) and HD individuals (Table S1). We found that RTP801 was enriched at the synaptosome subcellular compartment, along with synaptic markers SV2A and PSD-95. However, phospho-Ser473-Akt (P-Ser473-Akt) and Phospho-Ser235/236-S6 (P-Ser235/236-S6) were not significantly enriched in the crude synaptosomes (Figure S1). According to our previous results, we detected higher levels of RTP801 in homogenates derived from HD putamen (Figure S1A). Still, in homogenates, we confirmed that mTOR activities were elevated in HD as judged by increased levels of P-Ser473-Akt and P-Ser235/236-S6 (Figure S1A). Interestingly, we observed that RTP801 was significantly increased in HD synaptosomes (Figure 2B) in comparison to controls. As expected and in line with previous data^37^, postsynaptic marker PSD-95 was diminished in synaptosomes from HD patients and no differences were observed in pre-synaptic marker SV2A (Figure 2B). No significant differences were found in P-Ser473-Akt or P-Ser235/236-S6 at the synaptosomal compartment between non affected and HD individuals (Figure 2B).

**Figure 2.**
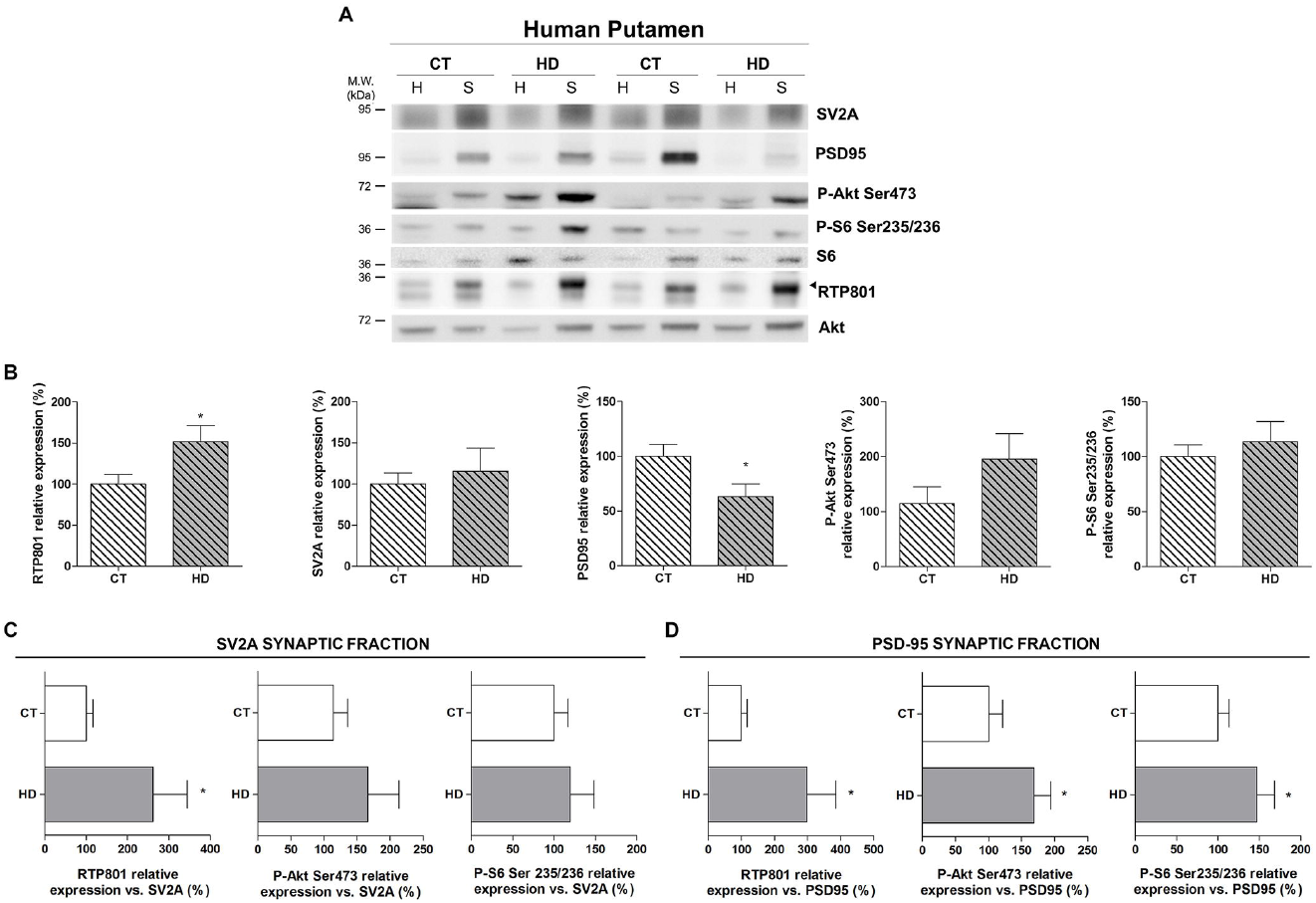
RTP801 is increased in the synaptic fraction derived from the putamen of HD patients. **(A,B)** Putamen of 6 HD patients and 5 control individuals lysates and synaptosomes were subjected to WB. Membranes were probed against RTP801, SV2A, PSD-95, P-Akt (Ser473), P-S6 (Ser235/236), total S6 and total Akt was used as a loading control. Graphs show the densitometric quantification of synaptosomal levels. **(C,D)** Graphs indicate the levels of RTP801, P-Akt (Ser473) and P-S6 (235/236) relative to synaptic markers (**C**) SV2A and (**D**) PSD-95 in the synaptosomes. The results are shown as mean ± SEM. Data were analyzed by Student’s T-test (\**P*<0.05).

In order to correct protein levels for the remaining synapses, synaptosomal levels of each protein were relativized to the levels of pre-synaptic marker SV2A (Figure 2C) and post-synaptic marker PSD-95 (Figure 2D). Hence, we confirmed that versus the SV2A- and PSD-95-synaptic proteins amount, RTP801 was increased in the HD group in comparison to controls. Furthermore, both P-Ser473-Akt and P-Ser235/236-S6 were elevated in the PSD-95-synaptic fraction of HD patients respect to unaffected individuals and showed the same tendency in the SV2A-synaptic fraction. These data are in line with the previously described impairment of the mTOR pathway in HD^38–40^.

Taken together, these results show that RTP801 is increased in the remaining synaptic terminals of HD patients’ putamen, along with the mTOR substrates P-Ser473-Akt and P-Ser235/236-S6.

To check whether these differences were region-specific, we performed the same analyses in the prefrontral cortex, a less affected brain region in the disease. Similarly, as in the putamen, RTP801, SV2A, PSD-95, P-Ser473-Akt and P-Ser235/236-S6 were highly enriched in the synaptosomal compartment (Figure S2, Enrichment), although no differences in RTP801 levels were observed neither in homogenates (Figure S2, Homogenates) nor in synaptosomes (Figure S3) between HD and non-affected individuals. Only synaptosomal P-Ser235/236-S6 normalized to SV2A or to PSD-95 showed a significant decrease in HD samples in contrast to control cases (Figure S3C&D).

### RTP801 is increased in striatal synapses of HD mouse models

To assess whether synaptic elevation of RTP801 could be reproduced in HD mouse models, we investigated the synaptic levels of RTP801 in the striatum of 10-month-old Hdh^Q7/Q111^ (KI) (Figure 3 & S4) and 16-week-old R6/1 mice (Figure 4 & S5). At this disease stage, these mice already display abnormal neuronal plasticity and motor and cognitive dysfunctions^41–46^.

**Figure 3.**
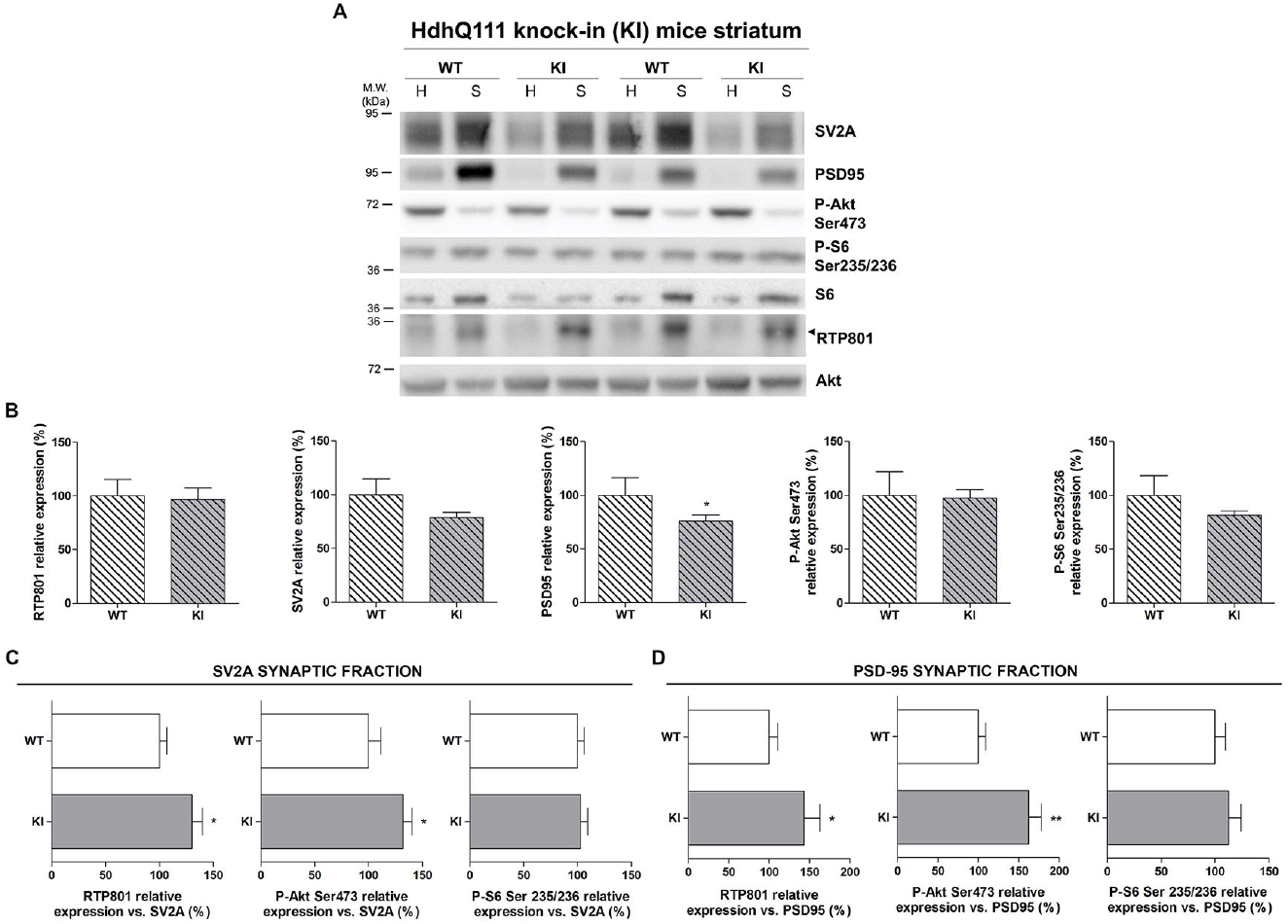
RTP801 is increased in the synaptic fraction derived from the striatum of HdhQ^7^/Q^111^ mice. **(A,B)** Striatal lysates and synaptosomes of 6 KI and 6 WT animals at 10-months of age were subjected to WB. Membranes were probed against RTP801, P-Akt (Ser473), P-S6 (Ser235/236), PSD-95, SV2A and total Akt was used as loading control. Graphs show the densitometric quantification of synaptosomal levels. **(C,D)** Graphs indicate the levels of RTP801, P-Akt (Ser473) and P-S6 (235/236) relative to synaptic markers (C) SV2A and (D) PSD-95 in the synaptosomes. The results are shown as mean ± SEM. Data were analysed by Student’s T-test (\**P*<0.05, \*\**P*<0.01).

**Figure 4.**
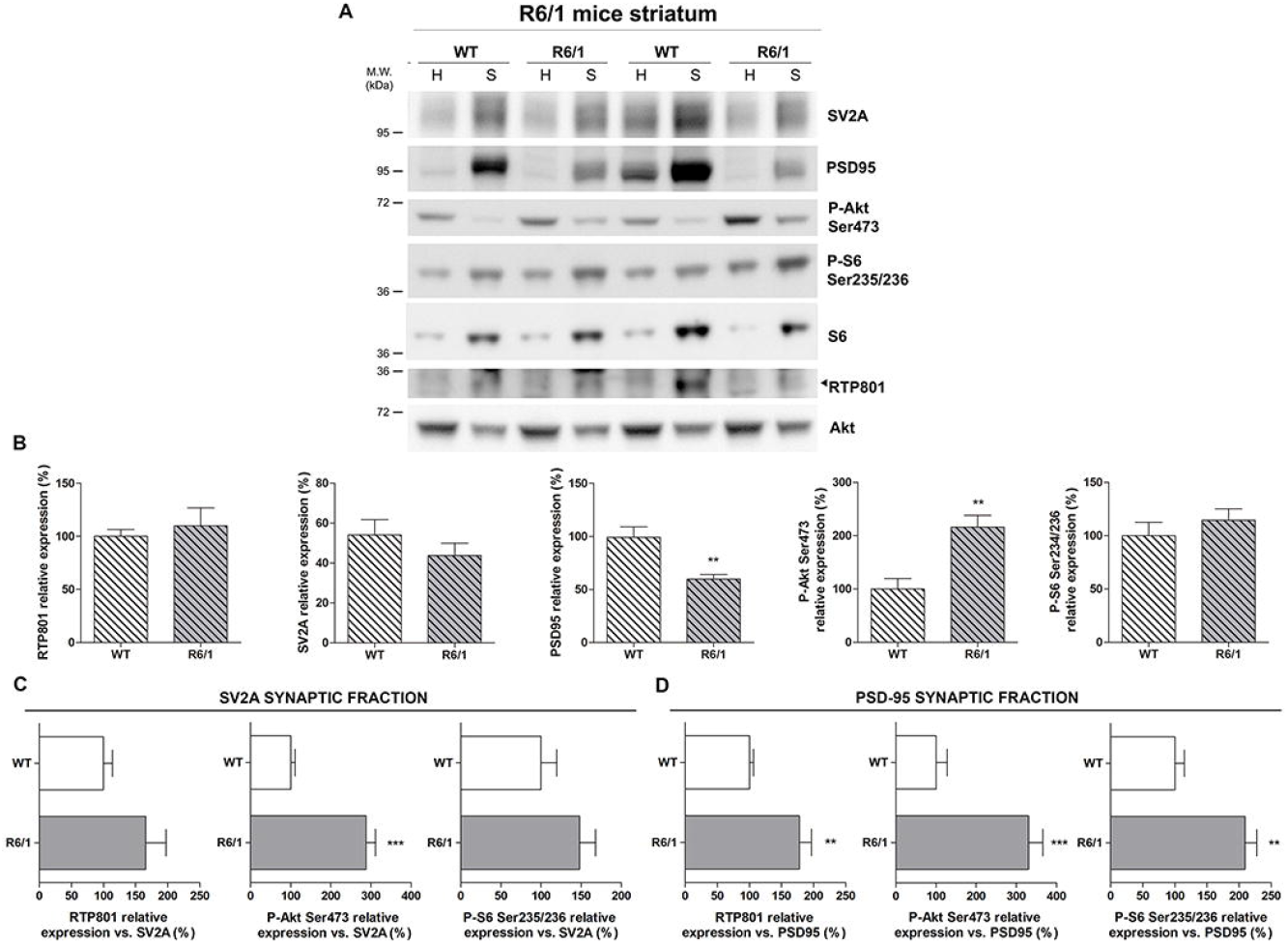
RTP801 is increased in the synaptic fraction derived from the striatum of R6/1 mice. Striatal lysates and synaptosomes of 7 R6/1 and 6 WT animals at 16-weeks of age were subjected to WB. Membranes were probed against RTP801, P-Akt (Ser473), P-S6 (Ser235/236), PSD-95, SV2A and total Akt as a loading control. Graphs show the densitometric quantification of synaptosomal levels. **(C,D)** Graphs indicate the levels of RTP801, P-Akt (Ser473) and P-S6 (235/236) relative to synaptic markers (C) SV2A and (D) PSD-95 in the synaptosomes. The results are shown as mean ± SEM. Data were analyzed by Student’s T-test (\*\**P*<0.01, \*\*\**P*<0.001).

Synaptosomal fractions of wild-type (WT) and HD mouse striata were enriched for RTP801, SV2A, PSD-95 and P-S6-Ser235/236 (Figure S4A-C, E; Figure S5A-C, E), as in human putamen. However, RTP801 protein levels did not differ between genotypes in either homogenates (Figure S4A and S5A) or synaptosomes (Figure 3B and 4B). PSD-95 displayed decreased protein levels in comparison to the WT group in KI and R6/1 mouse striatal synaptosomes (Figure 3B and 4B). After correcting the synaptic loss by expressing protein levels relative to the synaptic markers SV2A (Figure 3C & 4C) and PSD-95 (Figure 3D & 4D), RTP801 was increased in the synaptic fraction of KI and R6/1 mice. Interestingly, only P-Ser473-Akt was also significantly increased at the synaptic fraction of KI mice whereas in R6/1 mice, P-Ser235/236-S6 levels were also increased.

### RTP801 knockdown prevents motor learning deficits in symptomatic R6/1 mice

Since we found alterations in the striatal synaptic protein levels of RTP801 in HD mouse models and human HD brains, we investigated whether RTP801 could contribute to plasticity dysfunction in HD. To answer this question, 9-week-old WT and R6/1 mice were bilaterally injected with AAV-shCtr or AAV-shRTP801 into the striatum and 5 weeks later motor learning, a corticostriatal dependent function, was assessed by using the accelerating rotarod.

Comparable GFP signal was detected in the striatum of WT and R6/1 mice confirming the transduction of AAV-shCtr and AAV-shRTP801 (Figure 5A). Moreover, striatal injections of AAV-shRTP801 reduced RTP801 protein levels between 25-30%, both in WT and R6/1 mice (Figure 5B). The accelerating rotarod test showed that, as expected, shCtr-injected R6/1 animals performed poorly in this motor task in comparison to WT-shCtr-injected animals. Remarkably, RTP801 silencing in the R6/1 mice striatum prevented motor learning deficits as shown by the increased latency to fall, which was comparable to that in WT animals. Indeed, AAV-shCtr or AAV-shRTP801-injected WT mice showed no differences in motor learning skills (Figure 5C). These results indicate that mhtt-induced RTP801 synaptic increase is contributing to motor learning dysfunction in the R6/1 mouse model of HD.

**Figure 5.**
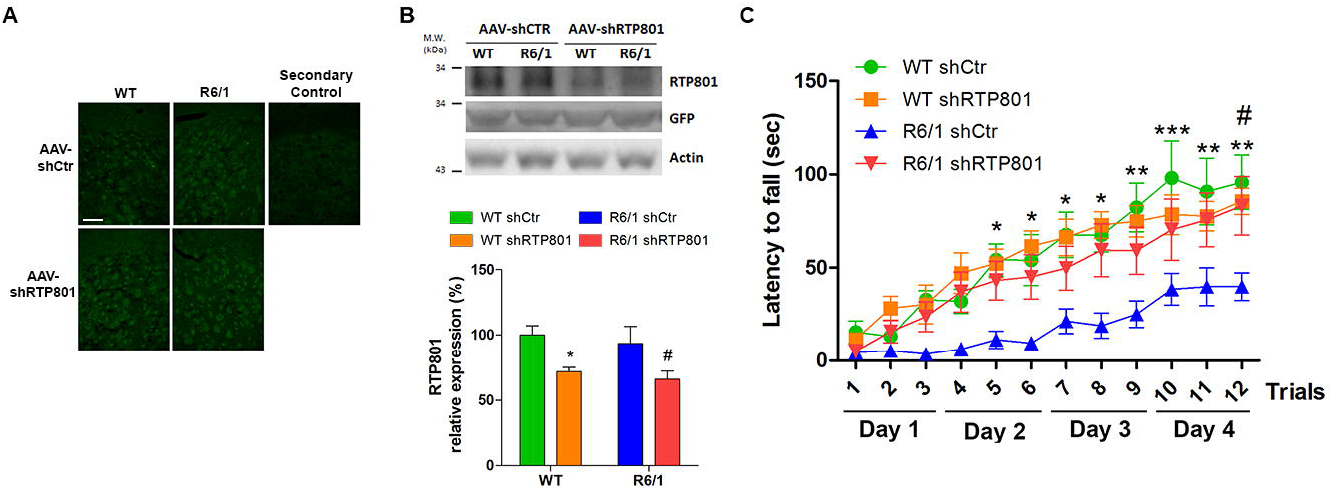
Striatal RTP801 knockdown preserves motor learning in symptomatic R6/1 mouse. **(A)** GFP immunofluorescence confirmed the transduction of AAV-shCtr and AAV-shRTP801 in WT and R6/1 mice striatal cells. A secondary control was performed without the incubation of primary antibody (anti-GFP). Scale bar 450μm. **(B)** Striatal lysates of WT and R6/1 injected with AAV-shCtr (n=6 WT and n=6 R6/1) or AAV-shRTP801 (n=6 WT and n=7 R6/1) were subjected to WB. Membranes were probed against RTP801, GFP and actin as a loading control. Graph show the densitometric quantification of RTP801 signal. Data are shown as a mean ± SEM and were analyzed with Two-way ANOVA followed by Bonferroni’s multiple comparisons test for *post-hoc* analyses (\**P*<0.05 *vs*. WT AAV-shCtr #*P*<0.05 *vs*. R6/1 AAV-shCtr). **(C)** Five weeks after adeno-associated transduction, motor learning was assessed in the same animals by the accelerating rotarod. The graph shows the latency to fall as the mean of three trials tested each day. Values are expressed as mean ± SEM and were analysed with Two-way ANOVA followed by Bonferroni’s multiple comparisons test for post-hoc analyses (\**P*<0.05, \*\**P*<0.01 and \*\*\**P*<0.001 R6/1 shCtr *vs*. WT-shCtr and #*P*<0.05 R6/1-shRTP801 *vs*. R6/1-shCtr).

### RTP801 knockdown normalizes phospho-Akt Ser473 and enhances the expression of synaptic proteins in the R6/1 mouse striatum

In order to understand the pathological function of synaptic RTP801 and how its knockdown prevents motor learning dysfunction, we analyzed by WB downstream effectors of the mTOR pathway in striatal synaptic fractions obtained from WT and R6/1 mice transduced with AAV-shCtr or AAV-shRTP801.

Knocking down RTP801 affected neither the levels of P-Ser2448-mTOR nor the levels of P-Ser235/236-S6, as mTORC1 readout (Figure 6A & 6B) but interestingly, it restored P-Ser473-Akt levels to basal levels both in homogenates and synaptosomes, with no significant effect in WT animals (Figure 6C). Therefore, this result indicates that RTP801 regulates the phosphorylation levels of Serine 473 Akt in this pathological context.

**Figure 6.**
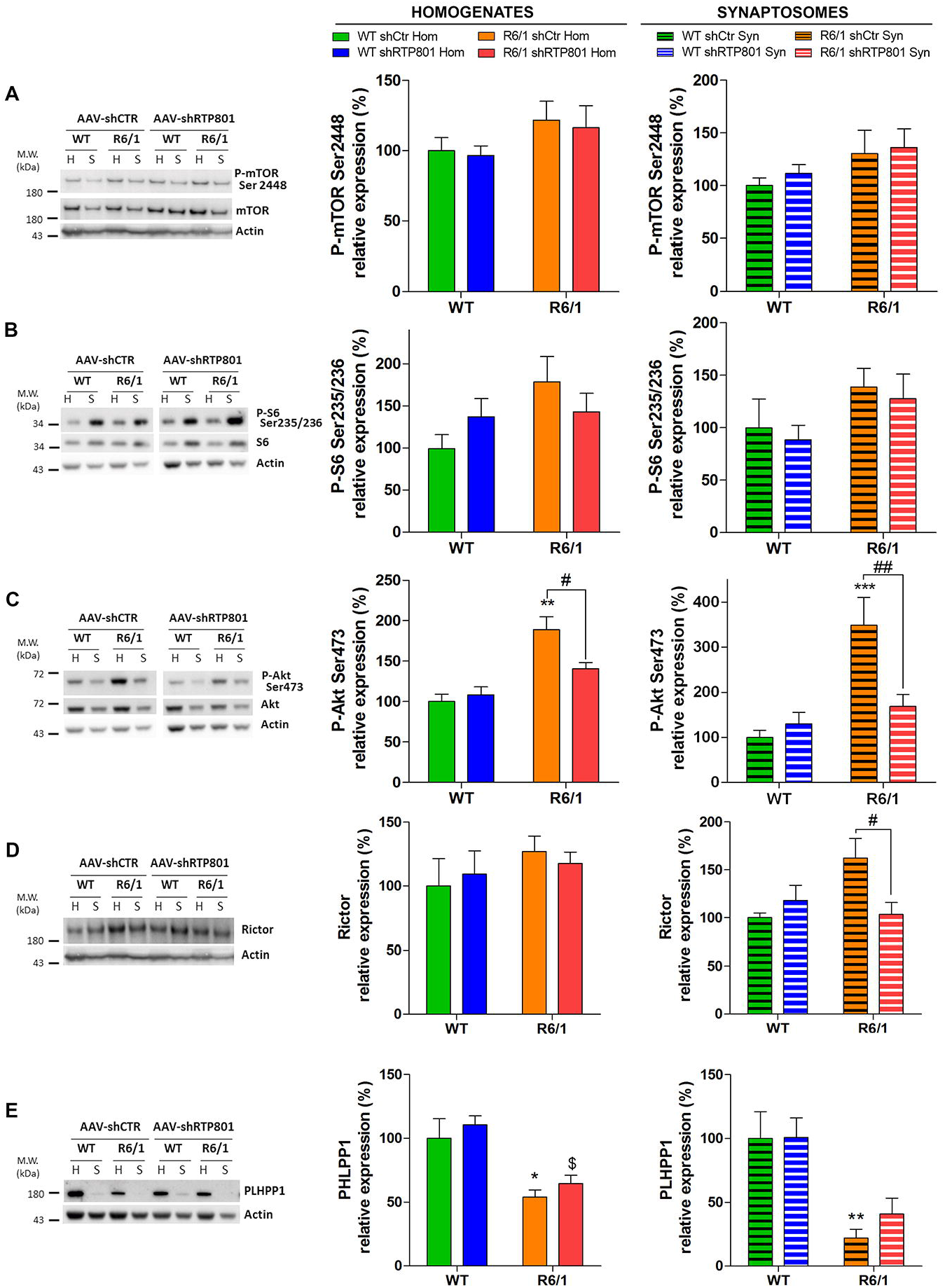
Knockdown of striatal RTP801 in R6/1 mice prevents hyperphosphorylation of Akt-(Ser473) by decreasing Rictor levels. Striatal lysates and synaptosomes of WT and R6/1 injected with AAV-shCtr (n=6 WT and n=6 R6/1) or AAV-shRTP801 (n=6 WT and n=7 R6/1) were subjected to WB. Membranes were probed against **(A)** P-mTOR (Ser2448), **(B)** P-S6 (Ser235/236), **(C)** P-Akt (Ser273), **(D)** Rictor and **(E)** PHLPP1. Total mTOR, S6, Akt and actin were used as loading controls. Graphs show the densitometric quantification. Values are shown as a mean ± SEM. Homogenates and synaptosomes data were analysed with Two-way ANOVA followed by Bonferroni’s multiple comparisons test for *post-hoc* analyses(\**P*<0.05, \*\**P*<0.01, \*\**P*<0.010 *vs*. WT AAV-shCtr; $P<0.05 *vs*. WT AAV-shRTP801; #*P*<0.05, ##*P*<0.01 *vs*. R6/1 AAV-shCtr).

Since increased levels of Rictor and reduced levels of PHLPP1 phosphatase are responsible for Akt hyperactivation in the R6/1 mouse model^23,24^, we investigated whether RTP801 knockdown in R6/1 mice could affect these two Akt modulators. Strikingly, RTP801 knockdown normalized specifically the synaptic levels of Rictor in R6/1 mice (Figure 6D) without altering the levels of PHLPP1 (Figure 6E). Hence, these results suggest that RTP801 knockdown prevents Akt hyperactivation by decreasing Rictor protein levels and thus downregulating the activity of mTORC2.

We next asked whether RTP801 downregulation could be translated to synaptic transmission modulation. Striatal RTP801 silencing induced an increase of AMPA receptor subunit GluA1 at the homogenate fraction in shRTP801-injected R6/1 mice in comparison to shCtr-injected R6/1 mice whereas non-significant tendencies were observed at the synaptic compartment (Fig 7A). One of the mechanisms contributing to synaptic impairment in the striatum of HD mouse models is the imbalance between BDNF receptors, TrkB and p75^NTR 47,48^. We found that RTP801 silencing increased TrkB levels in both homogenates and synaptosomes of R6/1 mice striatum in comparison to their shCtr-injected R6/1 littermates (Figure 7B) whereas p75^NTR^ was nor modified in any compartment analyzed (Figure 7C). Neither synaptosomal levels of PSD-95 (Figure 7D) nor NR1 (Figure 7E) were affected by the knockdown of RTP801.

**Figure 7.**
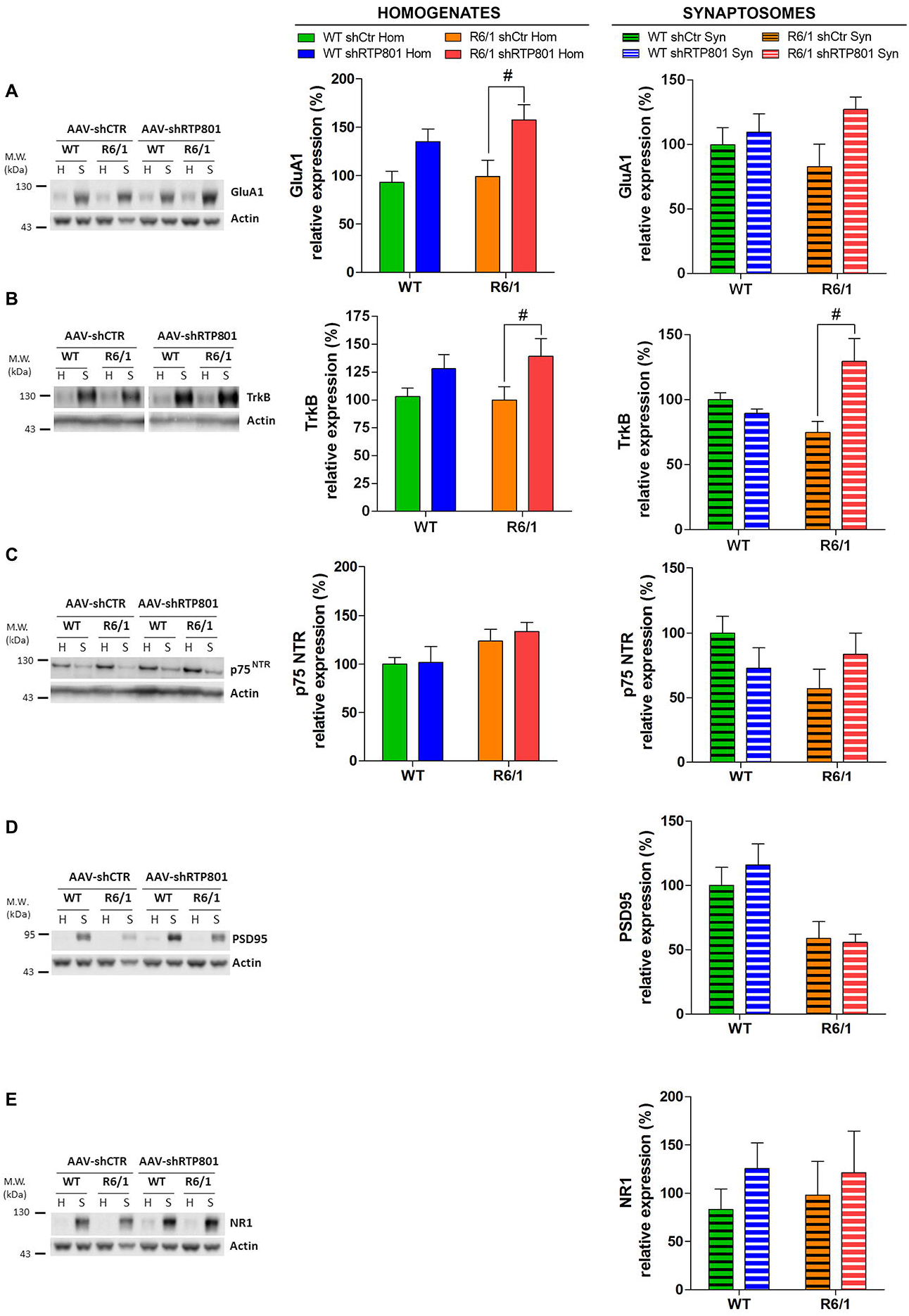
Knockdown of RTP801 in the striatum of R6/1 mice enhances the expression of synaptic proteins. Striatal lysates and synaptosomes of WT and R6/1 injected with AAV-shCtr (n=6 WT and n=6 R6/1) or AAV-shRTP801 (n=6 WT and n=7 R6/1) were subjected to WB. Membranes were probed against **(A)** GluA1, **(B)** TrkB, **(C)** p75^NTR^, **(D)** PSD-95 and **(E)** NR-1. Total actin was used as loading controls. Graphs show the densitometric quantification. Values are shown as a mean ± SEM. Homogenates and synaptosomes data were analysed with Two-way ANOVA followed by Bonferroni’s multiple comparisons test for *post-hoc* analyses; #*P*<0.05 *vs*. R6/1 AAV-shCtr).

These evidences indicate that striatal RTP801 knockdown prevents R6/1 motor learning deficit by decreasing Rictor levels, normalizing the bulk of P-Ser473-Akt and possibly, by enhancing postsynaptic signaling.

Altogether, the results indicate that RTP801 is increased in the synapses from the striatum, the most susceptible brain area to mhtt toxicity in both humans and HD mouse models, and its upregulation contributes to impair motor learning-related synaptic plasticity (Figure 8).

**Figure 8.**
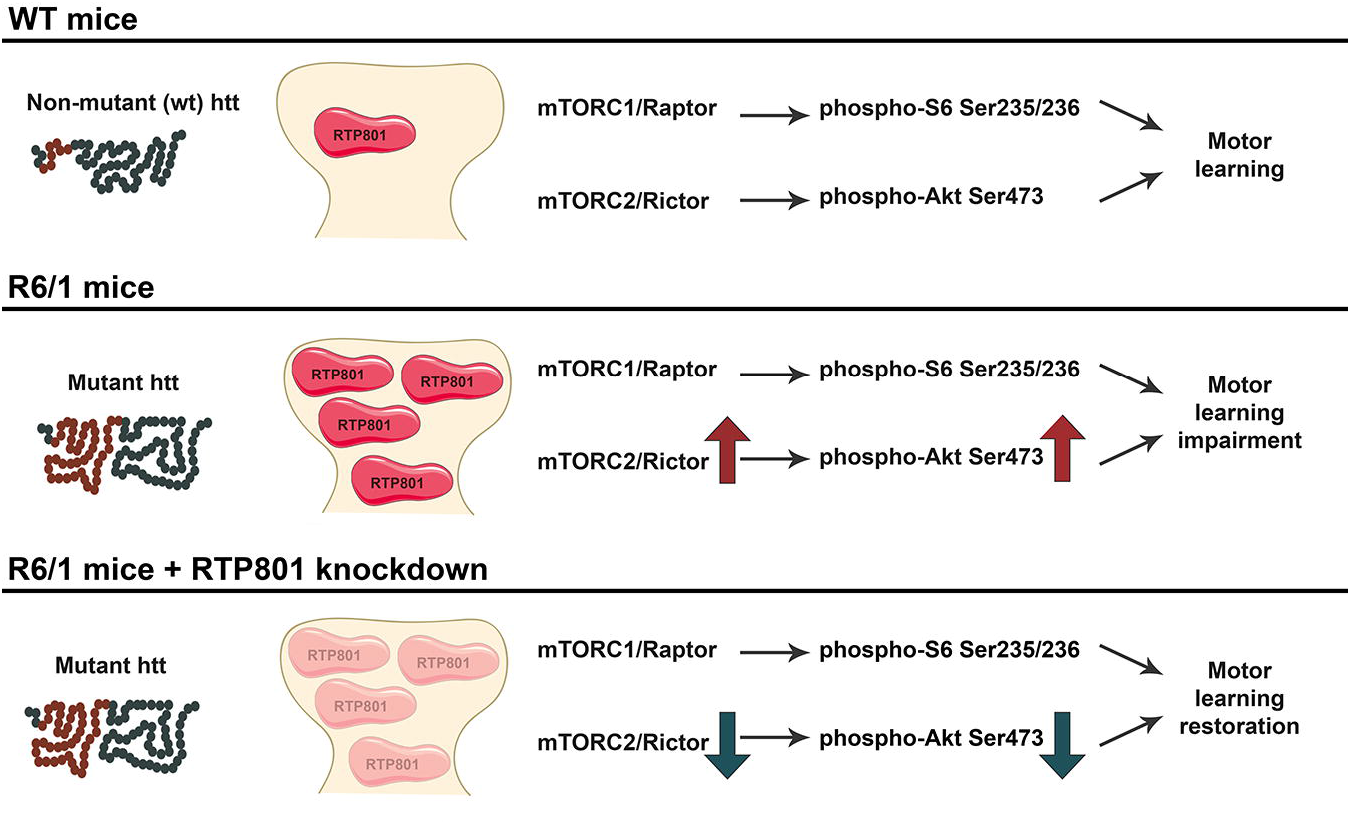
RTP801 knockdown prevents corticostriatal learning deficits. In comparison to WT mice, R6/1 mice display increased RTP801 at the striatal synapse along with increased Rictor and P-Akt (Ser473) and motor learning deficits. RTP801 silencing preserves corticostriatal motor learning function by decreasing the levels of Rictor and P-Akt Ser473 and also enhancing postsynaptic signaling by increasing GluA1 and TrkB receptors.

## DISCUSSION

HD is characterized by the appearance of aberrant synaptic plasticity and the progressive and selective loss of neuronal subpopulations^10,11,19,49–51^, in which mTOR plays a critical role^13^. In this work, we have investigated whether mhtt-induced RTP801 increase contributes to the impairment of synaptic plasticity in HD. Our results show that RTP801 is elevated in cultured neurons’ synapses overexpressing mhtt exon-1. Moreover, putamen, but not cortex, of HD patients displays increased levels of synaptic RTP801 respect to unaffected individuals. In addition, in HD mouse models, RTP801 is highly enriched in the striatal synaptic compartment when compared to WT animals. Notably, the knockdown of RTP801 expression in the striatum of R6/1 mice prevents motor learning deficits, which is accompanied by a synaptic restoration of Rictor and P-Ser473-Akt levels and an increase of GluA1 and TrkB receptors in the striatum.

We first demonstrated that mhtt elevates RTP801 protein in the soma and in the dendritic spines of primary cortical neurons, in accordance with our previous results obtained in differentiated PC12 cells^29^. Interestingly, mhtt-induced RTP801 up regulation courses along with a reduction in spine density. These results are in line with extensive work reporting the loss of dendritic spines in neurons overexpressing mhtt^36,52^ and suggest that RTP801 synaptic increase may be contributing to this phenomena.

Indeed, in human HD putamen, RTP801 protein levels were not only increased in whole lysates, as we previously described^29^, but also in synaptosomes. Structural and morphological synaptic alterations in striatal medium-sized spiny neurons^53,54^ and loss of dendritic spines in HD post-mortem brains have been widely reported^37,53,54^. In accordance with our findings, PSD-95 is also decreased in the striatum of HD patients^37^. Since we observed that RTP801 is enriched in the synaptic fraction and is highly accumulated in HD synaptic contacts in the striatum, we speculated that synaptic RTP801 in the remaining striatal dendritic spines could have a role in the altered plasticity associated to this neurodegenerative process. In addition, we found that RTP801 is also enriched in human cortical synapses, but no differences were found between controls and HD patients in the synaptic fraction. Even though morphological alterations in HD prefrontal cortical pyramidal neurons have been described^18^, our results indicate that RTP801 elevation is specific for the striatum. In line with this, neither the postsynaptic marker PSD-95 or the presynaptic marker SV2A were altered in the prefrontal cortex. Our results thus suggest that may be intrinsic neuronal properties modulate the effect of mhtt depending on the subcellular type.

In HD pathogenesis, synaptic impairment precedes motor symptoms^7–9^. In fact, HD mouse models are reported to exhibit cognitive deficits before the appearance of motor symptoms, indicating that abnormal synaptic plasticity occurs at early stages of the disease also in mouse models^46,55^.

In accordance with the lack of neuronal death in HD mouse models^41,42,56–58^, striatal levels of RTP801 in homogenates were not altered either in R6/1 or KI mice. However, specifically analyzing protein levels in synaptosomal preparations, we concluded that mhtt leads to a localized aberrant accumulation of RTP801 in synapses that could contribute to plasticity impairment in the pathology. In fact, structural alterations at the morphology and/or spine number in the striatum of the R6/1^59^ and the KI^60^ have been reported. This observation is also extended to other HD murine models such as N-terminal fragment R6/2 mice^61–63^ and transgenic full-length htt YAC128 mice^64^. As a synaptic marker, PSD-95 levels are reduced in those HD animal models^46,65,66^, supporting our results.

To evaluate the cortico-striatal function, responsible for motor learning^67^, R6/1 mice transduced with AAV-shCtr or AAV-shRTP801 were subjected to the accelerating rotarod test. Interestingly, silencing RTP801 in the striatum of R6/1 mice preserves their performance in the accelerating rotarod, comparable to WT mice, indicating that RTP801 knockdown retains the ability to learn new motor skills in the HD mice. The fact that silencing RTP801 in WT animals does not affect their performance at this paradigm could suggest that remaining basal levels of RTP801 are sufficient to maintain proper motor learning plasticity and, therefore, cortico-striatal functionality. Highlighting the function of RTP801 in synaptic plasticity, its downregulation in the Substantia Nigra pars compacta has also been shown to restore motor learning skills in a PD mice model subjected to chronic stress^68^.

The only known function of RTP801 is as a mTOR signaling repressor. In PD studies using cellular and animal models, RTP801 upregulation upon cell stress represses mTOR and finally inhibits Akt pro-survival signals, triggering neuronal death. In PD cellular models and nigral neurons of PD brains, RTP801 increases along with a decrease in the phosphorylation of Akt and S6^30,31^. Indeed, in HD, alterations in the activity of both mTORC1^40^ and mTORC2^24^ have been described. Here, we confirm that both mTOR complexes display aberrant activity and, importantly, that alterations found at total cell level are extended to the synaptic compartment.

Unlike in PD models, we show that mhtt increases RTP801 despite the hyperphosphorylation of Akt at Ser473 residue in the synaptosomes. Interestingly, P-Ser235/236-S6 displays the same trend. The fact that RTP801 levels are increased in HD models may also be a consequence of mTOR pathway hyperactivity, since RTP801 protein synthesis is mTORC1-dependent^69^.

Supporting its synaptic role, RTP801 silencing also alters the levels of mTOR pathway-associated protein components and enhances the expression of postsynaptic proteins. Our data indicate that RTP801 downregulation in R6/1 mice striatum prevents Akt hyperphosphorylation at Ser473 residue, both at homogenates and synaptosomes. Overactivation of Akt has been proposed to be a pro-survival response to counteract mhtt toxicity in the disease^22–24^ However, some evidences have also pointed out that synaptic plasticity alterations occur along with hyperphosphorylation of Akt^70^ and that sustained overactivation of Akt could be detrimental for cell survival and synaptic function^71,72^. Moreover, the function of mTOR complexes can differ between neuronal compartments. For instance, mTORC2 at the soma mediates the activation of Akt to ensure neuronal survival, whereas mTORC2 close to the synapse mediates the activation of PKC, responsible for the polymerization of actin and receptor trafficking (reviewed in^73^. Thus, RTP801 function could also be different between cell compartments. Altogether, this evidence confirms the regulation of RTP801 over mTORC2-dependent Akt phosphorylation but, in the presence of mhtt, the regulation is not negative.

Phosphorylation levels of Akt at Ser473 depend on a proper activity of the mTOR kinase complexed with Rictor^74^ and the phosphatase PHLPP1^25,26^, that specifically dephosphorylates this residue. In fact, in HD, increased Rictor and decreased PHLPP1 levels are both responsible for Akt overactivation^23,24^. In line with that, we found that Rictor synaptic up regulation in the R6/1 mice is sensitive to RTP801 levels since RTP801 knockdown specifically reduces synaptosomal Rictor levels without affecting PHLPP1. Still the regulation of RTP801 towards Rictor or whether it exists any feedback mechanisms over Akt phosphorylation remains uncertain.

Interestingly, we observed that RTP801 knockdown in the striatum elevated the levels of AMPA subunit receptor GluA1. AMPA receptors normally are heterotetramers composed mostly by calcium impermeable GluA2 subunits. However, GluA1 subunit confers calcium permeability to the pore, allowing signaling activation and synaptic regulation^75–77^. Therefore, we speculate that this GluA1 up regulation could affect the composition of the AMPA receptors and promote synaptic strength. In line with this, RTP801 knockdown elevated TrkB levels in R6/1 homogenates and the same trend was observed in the synaptosomal fraction, with no changes in p75^NTR^. Both TrkB and p75^NTR^ are BDNF receptors, although each receptor activates different signaling pathways. BDNF mediates neuronal survival by the activation of TrkB receptor^78,79^ whereas p75^NTR^ can activate apoptotic signals leading to neuronal death^47,80–83^. In fact, imbalance of TrkB and p75^NTR^ has been proposed to mediate impaired plasticity in HD^47,84^. Hence, increased TrkB and GluA1 levels, as a result of RTP801 silencing, could promote neuroprotection and enhancement of synaptic transmission^78,79,84^.

## CONCLUSIONS

Our results indicate that downregulation of striatal RTP801 prevents motor learning deficits in R6/1 mouse model, in part by by preventing Akt hyperphosphorylation at the Ser473 residue via Rictor downregulation and by increasing postsynaptic GluA1 and TrkB receptors. Hence, RTP801 emerges as a promising target for future therapeutic strategies to prevent or at least halt the progression of HD.

## METHODS

### Cortical primary neurons

Primary cortical cultures were obtained from rat Sprague-Dawley embryos at day 18 (E18) as previously described^29^. Briefly, cortices were dissected out, dissociated in 0.05% trypsin (Sigma-Aldrich) and maintained in Neurobasal medium supplemented with B27 and 2mM GlutaMAX™ (all from Thermo Fisher Scientific) in a 5% CO_2_ humified atmosphere at 37°C. For imaging experiments, neurons were plated at a density of 300 cells/mm^2^ on poly-L-lysine-coated (Sigma-Aldrich) in seeding medium containing MEM medium (Thermo Fisher Scientific) supplemented with 10% heat-inactivated horse serum (Sigma-Aldrich). After 60 minutes, medium was replaced for supplemented Neurobasal. Medium was replenished one third every 7 days.

Transfection of the constructs eGFP, Q25, Q72 and Q103 and immunocytochemistry of neuronal cultures were performed as described elsewhere^85^. Briefly, cultures were incubated with the primary antibody mouse monoclonal against eGFP (1:1000, Santa Cruz Biotechnology) and rabbit polyclonal against RTP801 (1:150, Proteintech Group Inc.). AlexaFlour Phalloidin 568 (1:1000, Thermo Fisher Scientific) was diluted along with the secondary antibody during incubation. The secondary antibodies used were goat anti-Mouse IgG (H+L) conjugated to AlexaFluor™ 488 or goat anti-Rabbit IgG (H+L) conjugated with Alexa Fluor™ 568 or AlexaFluor™ 647 (all from Thermo Fisher Scientific). For nuclear staining, nuclei were revealed with Hoechst 3342 (Thermo Fisher Scientific) diluted 1:5000 along with the secondary antibody incubation. In the case of actin filamentous labelling, AlexaFluor™ 568 Phalloidin diluted 1:10000 was incubated along with the secondary antibody and Hoechst 3342. Spine density was calculated by scoring the number of spines for each 15 μm dendrite-length of at least 10 dendrites per condition, in three independent experiments. ImageJ was used to quantify RTP801 intensity as the Integrated Density in the soma and in the dendrites.

### HD mouse models

R6/1 transgenic mice expressing the exon-1 of mhtt with 145 CAG repeats were obtained from Jackson Laboratory and maintained in B6CBA background. Hdh^Q7^ WT mice, with 7 CAG repeats, and Hdh^Q111^ knock-in mice, with targeted insertion of 109 CAG repeats that extends the glutamine segment in murine htt to 111 residues, were maintained on a C57Bl/6 genetic background. Male and female Hdh^Q7/Q111^ heterozygous were intercrossed to generate age-matched Hdh^Q7/Q7^ WT and Hdh^Q7/Q111^ knock-in littermates. All mice used were males and were housed together in numerical birth order in groups of mixed genotypes. Data were recorded for analysis by microchip mouse number. Animals were housed with access to food and water ad libitum in a colony room kept at 19-22°C and 40-60% humidity, under a 12:12 hours light/dark cycle. All procedures were carried out in accordance with the National Institutes of Health Guide for the Care and Use of Laboratory Animals and approved by the local animal care committee of the Universitat de Barcelona, following European (2010/63/UE) and Spanish (RD53/2013) regulations for the care and use of laboratory animals.

### Human postmortem tissue

Postmortem striatal and frontal cortical brain tissues from control and HD patients were obtained from the Neurological Tissue Bank (Biobank-HC-IDIBAPS) thanks to Dr. Ellen Gelpi collaboration. (See **Supplementary Table 1**)

### Crude synaptosomal preparations

Synaptosomes were prepared from mouse striata and human putamen and frontal cortex as previously described elsewhere with minor modifications. Briefly, dissected tissue was homogenized in Krebs-Ringer (KR) buffer (125 mM Nacl, 1.2 mM KCl, 22 mM NaHCO_3_, 1 mM HaH_2_PO_4_, 1.2 mM MgSO_4_, 1.2 mM CaCl_2_, 10 mM glucose (pH=7.4)) supplemented with 0.32 M Sucrose. Then, samples were subjected to a first centrifugation at 1,000g for 10 minutes (4°C) to discard debris. A sample of the supernatant was kept as the homogenate fraction. After, the supernatant was subjected to a second centrifugation at 16,000g for 15 minutes (4°C) to obtain the crude synaptosomal fraction. The pellet was finally resuspended in KR 0.32M sucrose buffer and subjected to WB to analyze protein content.

### Western blotting

Homogenate and synaptosomal protein analysis was performed by Western blotting as previously described^86^. The following primary antibodies were used: Akt rabbit polyclonal (1:1000, Cell Signaling), Akt-phospho Ser473 rabbit polyclonal (1:1000, Cell Signaling), mTOR rabbit polyclonal (1:1000, Cell Signaling), mTOR-phospho Ser2448 (1:1000, Cell Signaling), anti-NR1 mouse monoclonal (1:1000, Chemicon), p75^NTR^ rabbit polyclonal (1:1000, Thermo Fisher Scientific), PHLPP1 rabbit polyclonal (1:500, Cayman Chemical), PSD-95 mouse monoclonal (1:1000, Thermo Fisher Scientific), RTP801 (1:500, Proteintech Group Inc.), RPS6-phospho Ser235/236 rabbit polyclonal (1:1000, Cell Signaling), SV2A mouse monoclonal (1:1000, Santa Cruz Biotechnology) and TrkB mouse monoclonal (1:1000, BD Bioscience). Then, membranes were incubated with the corresponding secondary antibodies: goat anti-Mouse IgG (H+L) and goat anti-Rabbit IgG (H+L) conjugated to horseradish peroxidase (1:10,000, Thermo Fisher Scientific). Loading control was obtained incubating with anti-actin-horseradish peroxidase conjugated (1:100,000, Sigma Aldrich). Chemiluminescent images were acquired using a LAS-3000 imager (Fuji) or ChemiDoc (Bio-Rad) imaging systems and quantified by computer-assisted densitometric analysis (Image J software).

### Intrastriatal injection of adeno-associated vectors

To knockdown RTP801 expression, shRNA verified scrambled sequence (5’-GTGCGTTGCTAGTACCAAC-3’) and the one against RTP801 (5’-AAGACTCCTCATACCTGGATG-3’) were cloned into a rAAV2/8-GFP adenoviral vector at restriction sites BamHI at 5’ and Agel at the 3’. The rAAV2/8 plasmids and infectious AAV viral particles containing GFP expression cassette with shCtr or shRTP801 were generated by the Vectors Production Unit from the Center of Animal Biotechnology and Gene Therapy at the

Universitat Autònoma de Barcelona. Nine-week-old WT and R6/1 mice were subjected to bilateral intrastriatal injections of rAVV2/8 expressing shRTP801 or control shRNA. Animals were deeply anesthetized with a mixture of oxygen and isoflurane (4-5 for induction and 1-2 for maintaining anesthesia) and placed in a stereotaxic apparatus for bilateral intrastriatal injections (2 μl; 15□×□10^9^ genomic copies). Two injections were performed in the striatum at the following coordinates relative to bregma: (1) anteroposterior (AP), +□0.8; mediolateral (ML), +/-□1.8; and profundity, −2.6 mm and (2) AP, +□0.3; ML, +/- □2; and profundity, −2.6 mm, below the dural surface. Viral vectors were injected using a 10 μl-Hamilton microliter syringe at an infusion rate of 200 nl/min. The needle was left in place for 5 min to ensure complete diffusion of the viruses and then slowly retracted from the brain. Both hemispheres were injected with the same shRNA.

### Accelerating rotarod

Five weeks after intrastriatal injection of rAAV2/8-shRNAs, animals were subjected to the accelerating rotarod test to analyze motor learning. Mice were placed on a 3 cm rod (Panlab) with an increasing speed from 4 to 40 rpm over 5 minutes. Latency to fall was recorded as the time mice spent in the rod before falling. Accelerating rotarod test was performed for 4 days, 3 trials per day. Trials in the same day were separated by 1 hour.

### Immunohistofluorescence of mouse brain sections

Animals were deeply anesthetized with dolethal (Vétoquinol) (60 mg/kg), and intracardially perfused with 4% paraformaldehyde in PBS buffer (pH=7.2-7.4). Brains were removed and post-fixed for 18-24h, and cryoprotected with 30% sucrose in PBS 0.02% sodium azide and frozen in dry-ice cooled 2-methylbutane. Serial cryostat 25 μm-thick sections were collected in PBS 0.2% sodium azide as free-floating sections and processed for immunohistofluorescence. Sections were washed with PBS and incubated with 50 mM NH_4_Cl for 30 minutes to block aldehyde free-induced fluorescence. Tissue was blocked with SuperBlock-PBS containing 0.3% Triton X-100 for 2 hours at room temperature. After, slices were incubated overnight at 4°C with the primary antibodies anti-GFP chicken polyclonal (Synaptic Systems) diluted 1:500 in SuperBlock-PBS 0,3% Triton X-100. Later, sections were washed in PBS and incubated for 2 hours at room temperature with the secondary antibody goat anti-Chicken IgY (H+L) conjugated to AlexaFluor™ 488 diluted 1:500 in SuperBlock-PBS 0.3% Triton X-100. For nuclear staining, nuclei were revealed with Hoechst 3342 diluted 1:5000 along with the secondary antibody incubation. Slices were washed with PBS before being mounted with ProLong™ Gold Antifade Mountant on SuperFrost™ Plus Adhesion Slides (Thermo Fisher Scientific). Stained mouse brain sections were observed by epifluorescent microscopy.

### Statistical analysis

All experiments were performed at least in triplicate, and results are reported as mean+SEM. Student’s T-test was performed as unpaired, two-tailed sets of arrays and presented as probability *P* values. One-way ANOVA with Bonferroni’s multiple comparison test and Two-way ANOVA followed by Bonferroni’s post-hoc tests were performed for the comparison of multiple groups. Values of *P* < 0.05 were considered as statistically significant.

## ABBREVIATIONS

(HD): Huntington’s disease
(KO): knock out
(mTOR): mechanistic target of rapamycin
(mTORC): mTOR complex
(mhtt): mutant huntingtin
(PSD): postsynaptic density
(WB): Western blot
(WT): wild type

## ACKNOWLEDGEMENTS

The authors thank Dr. M MacDonald (Massachusetts General Hospital, Boston, Massachusetts, USA) for the Hdh^Q7/Q111^ mice. The constructs Q25 Q72 and Q103 were a kind gift of Dr. G.M. Lawless (Cure HD Initiative, Reagent Resource Bank of the Hereditary Disease Foundation, New York, NY). We also thank the Neurological Tissue Bank of the Biobanc-Hospital Clínic-IDIBAPS (Barcelona, Spain) and Dr. Ellen Gelpi for providing human tissue samples, and Ana López for her technical support. We thank Maria Calvo from the Advanced Microscopy Unit, Scientific and Technological Centers, University of Barcelona, for their support and advice in confocal techniques.

## DECLARATIONS

### Declarations

#### Ethical Approval and Consent to participate

All procedures were performed in compliance with the NIH Guide for the Care and Use of Laboratory Animals and approved by the local animal care committee of Universitat de Barcelona following European (2010/63/UE) and Spanish (RD53/2013) regulations for the care and use of laboratory animals.

Human samples were obtained following the guidelines and approval of the local ethics committee (Hospital Clínic of Barcelona’s Clinical Research Ethics Committee).

#### Consent for publication

Non applicable

#### Availability of supporting data

All data generated or analyzed during this study are included in this published article and the supplementary information files.

#### Authors’ contributions

N.M-F., L.P-S., J.C-M., M.M., S.G., J.A., E.P-N. and C.M. have contributed in the conception and design of the study, acquisition and analysis of data and in drafting the manuscript and figures. All authors read and approved the final manuscript.

## SUPPLEMENTAL INFORMATION TITLES AND LEGENDS

**Figure S1. Synaptosomal enrichment of proteins in the putamen of HD patients**. Putaminal lysates and synaptosomes of 5 HD patients and 5 control individuals were subjected to WB. Membranes were probed against **(A)** RTP801, **(B)** SV2A, **(C)** PSD-95, **(D)** P-Akt (Ser473) and **(E)** P-S6 (Ser235/236) and total Akt as a loading control. Graphs show the densitometric quantification. Data is shown as a mean ± SEM. Data were analyzed with Two-way ANOVA followed by Bonferroni’s multiple comparisons test for post-hoc analyses (\*\*\**P*<0.001 *vs*. CT homogenate) and data of homogenates and synaptosomes were analyzed by Student’s T-test (\**P*<0.05, \*\**P*<0.01).

**Figure S2. Synaptosomal enrichment of proteins in the cortex of HD patients**. Prefrontal cortex lysates and synaptosomes of 5 HD patients and 6 control individuals were subjected to WB. Membranes were probed against **(A)** RTP801, **(B)** SV2A, **(C)** PSD-95, **(D)** P-Akt (Ser473) and **(E)** P-S6 (Ser235/236) and total Akt as a loading control. Graphs show the densitometric quantification. Data is shown as a mean ± SEM. Enrichment data was analyzed with Two-way ANOVA followed by Bonferroni’s multiple comparisons test for post-hoc analyses (\**P*<0.05, \*\**P*<0.01, \*\*\**P*<0.001 *vs*. CT homogenate; ### *P*<0.001 *vs*. HD homogenate). Data from homogenates was analyzed with Student’s t-test.

**Figure S3. RTP801 is not altered in the synaptic fraction derived from the cortex of HD patients. (A, B)** Prefrontal cortex lysates and synaptosomes of 6 HD patients and 6 control individuals were subjected to WB. Membranes were probed against RTP801, P-Akt (Ser473), P-S6 (Ser235/236), PSD-95, SV2A and total Akt as a loading control. Graphs show the densitometric quantification of synaptosomal levels. **(C, D)** Graphs indicate the levels of RTP801, P-Akt (Ser473) and P-S6 (235/236) relative to synaptic markers **(C)** SV2A and **(D)** PSD-95 in the synaptosomes. The results are shown as mean ± SEM. Data were analyzed by Student’s T-test (\**P*<0.05, \*\**P*<0.01).

**Figure S4. Synaptosomal enrichment of proteins in the in the striatum of HdhQ^7^/Q^111^ mice**. Striatal lysates and synaptosomes of 6 KI and 6 WT animals at 10-months of age were subjected to WB. Membranes were probed against **(A)** RTP801, **(B)** SV2A, **(C)** PSD-95, **(D)** P-Akt (Ser473) and **(E)** P-S6 (Ser235/236) and total Akt as a loading control. Graphs show the densitometric quantification. Data is shown as a mean ± SEM. Data were analyzed with Two-way ANOVA followed by Bonferroni’s multiple comparisons test for post-hoc analyses (\*\**P*<0.01, \*\*\**P*<0.001 *vs*. WT homogenate, ##*P*<0.01, ###*P*<0.001 *vs*. KI homogenate) and data of homogenates were analysed by Student’s T-test (\**P*<0.05).

**Figure S5. Synaptosomal enrichment of proteins in the striatum of R6/1 mice**. Striatal lysates and synaptosomes of 7 R6/1 and 6 WT animals at 16-weeks of age were subjected to WB. Membranes were probed against **(A)** RTP801, **(B)** SV2A, **(C)** PSD-95, **(D)** P-Akt (Ser473) and **(E)** P-S6 (Ser235/236) and total Akt as a loading control. Graphs show the densitometric quantification. Data in shown as a mean ± SEM. Data were analysed with Two-way ANOVA followed by Bonferroni’s multiple comparisons test for post-hoc analyses (\**P*<0.05, \*\**P*<0.01, \*\*\**P*<0.001 *vs*. WT homogenate, #*P*<0.05, ###*P*<0.001 *vs*. R6/1 homogenate) and data of homogenates were analysed by Student’s T-test (\*\*\**P*<0.001).

**Figure S6. Enrichment of proteins in the striatum of R6/1 after RTP801 knockdown**. Striatal lysates and synaptosomes of WT and R6/1 injected with *AAV-shCtr* (n=6 WT and n=6 R6/1) or AAV-shRTP801 (n=6 WT and n=7 R6/1) were subjected to WB. Graphs show the densitometric quantification of **(A)** P-mTOR (Ser2448), **(B)** P-S6 (Ser235/236), **(C)** P-Akt (Ser273), **(D)** Rictor, **(E)** PHLPP1, **(F)** GluA1, **(G)** TrkB and **(H)** p75^NTR^. Values are shown as a mean ± SEM. Enrichment data were analyzed with Two-way ANOVA followed by Bonferroni’s multiple comparisons test for *post-hoc* analyses (\*\**P*<0.01, \*\*\**P*<0.001 *vs*. WT AAV-shCtr Hom; #*P*<0.05, ##*P*<0.01, ###*P*<0.001 *vs*. R6/1 AAV-shCtr Hom;; ++*P*<0.01, +++*P*<0.001 *vs*. WT AAV-shRTP801; $*P*<0.05, $$*P*<0.01, $$$*P*<0.001 *vs*. R6/1-shRTP801 Hom).

