## Supplementary figures and images for "Synaptic RTP801 Contributes to Motor Learning Dysfunction in Huntington’s Disease"

### Supplementary material

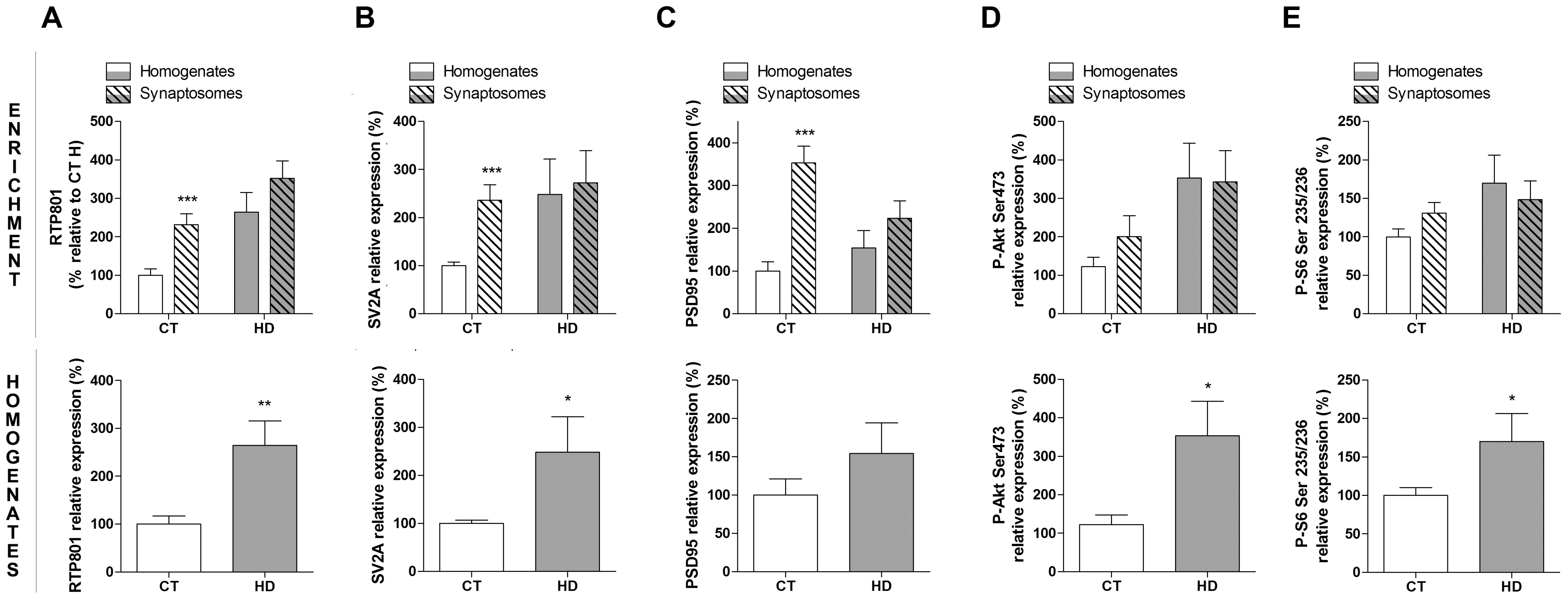

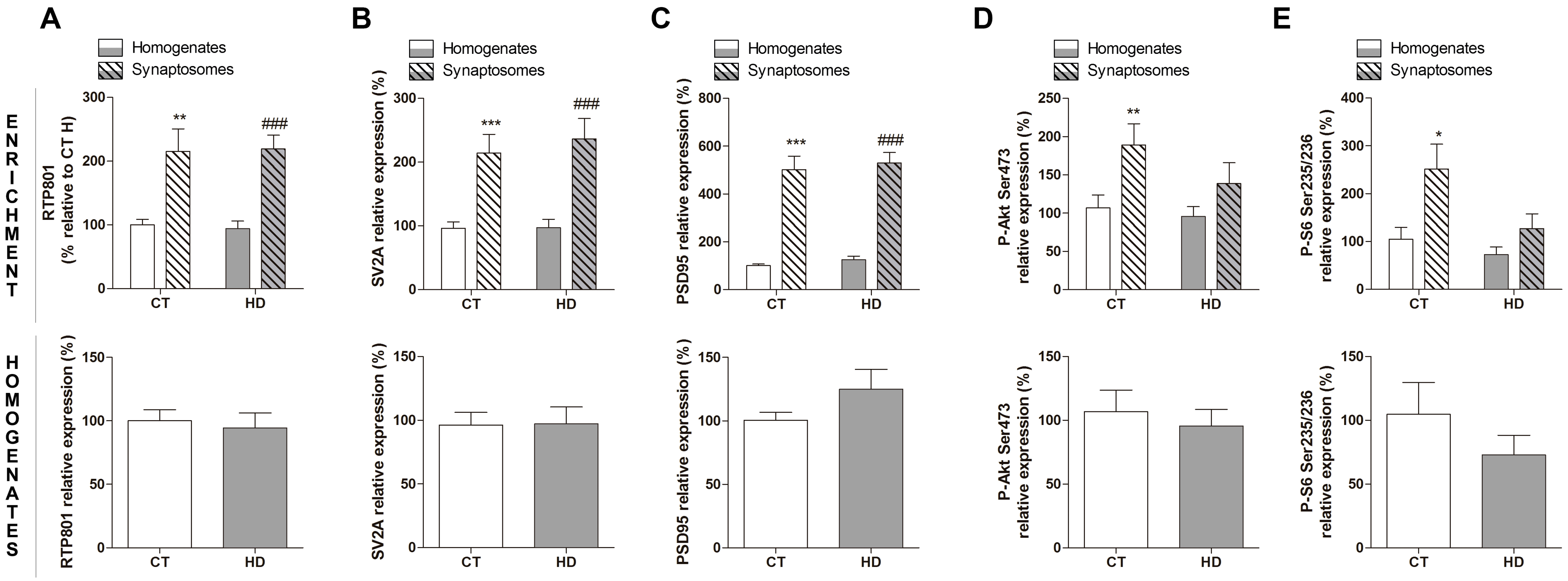

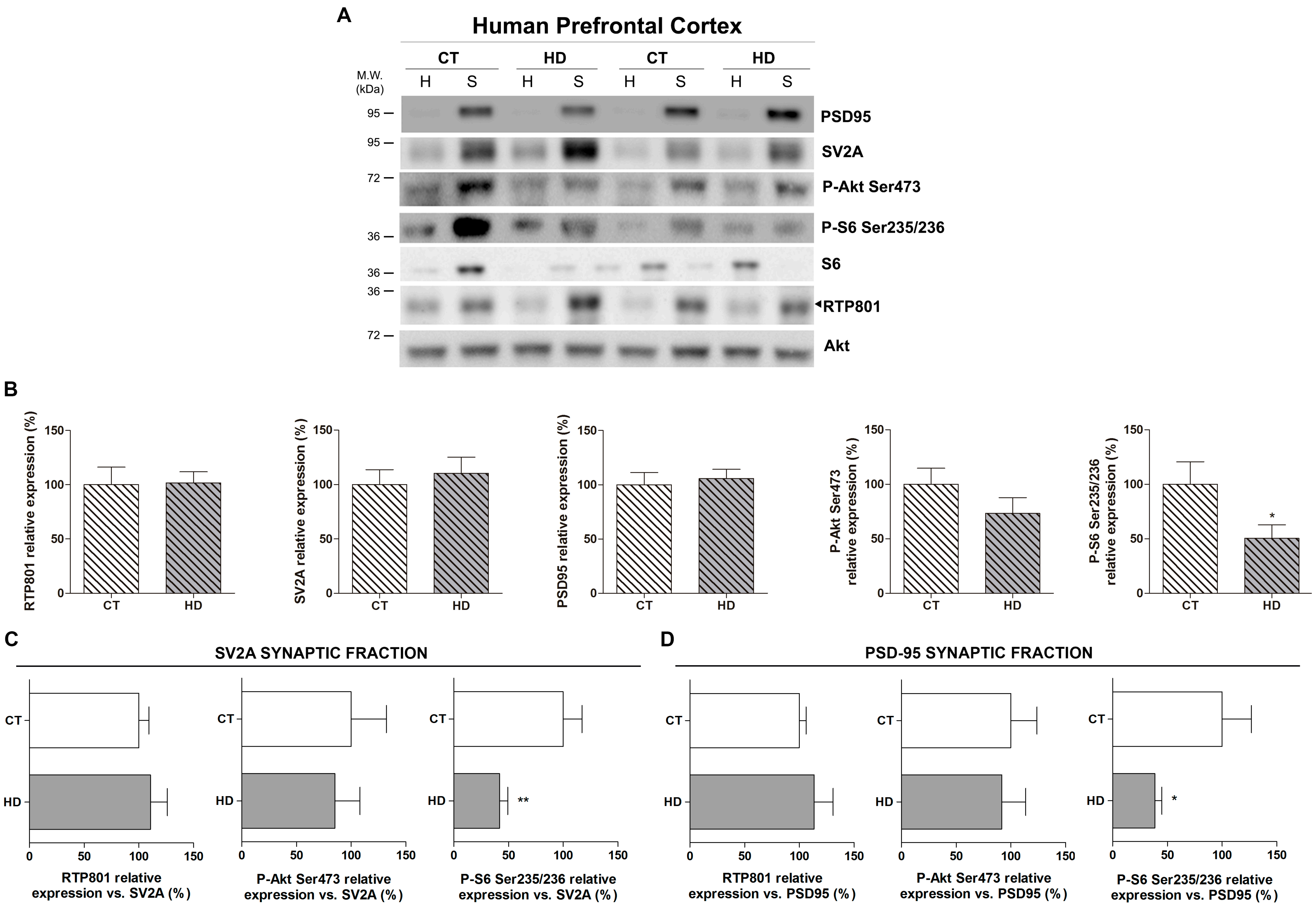

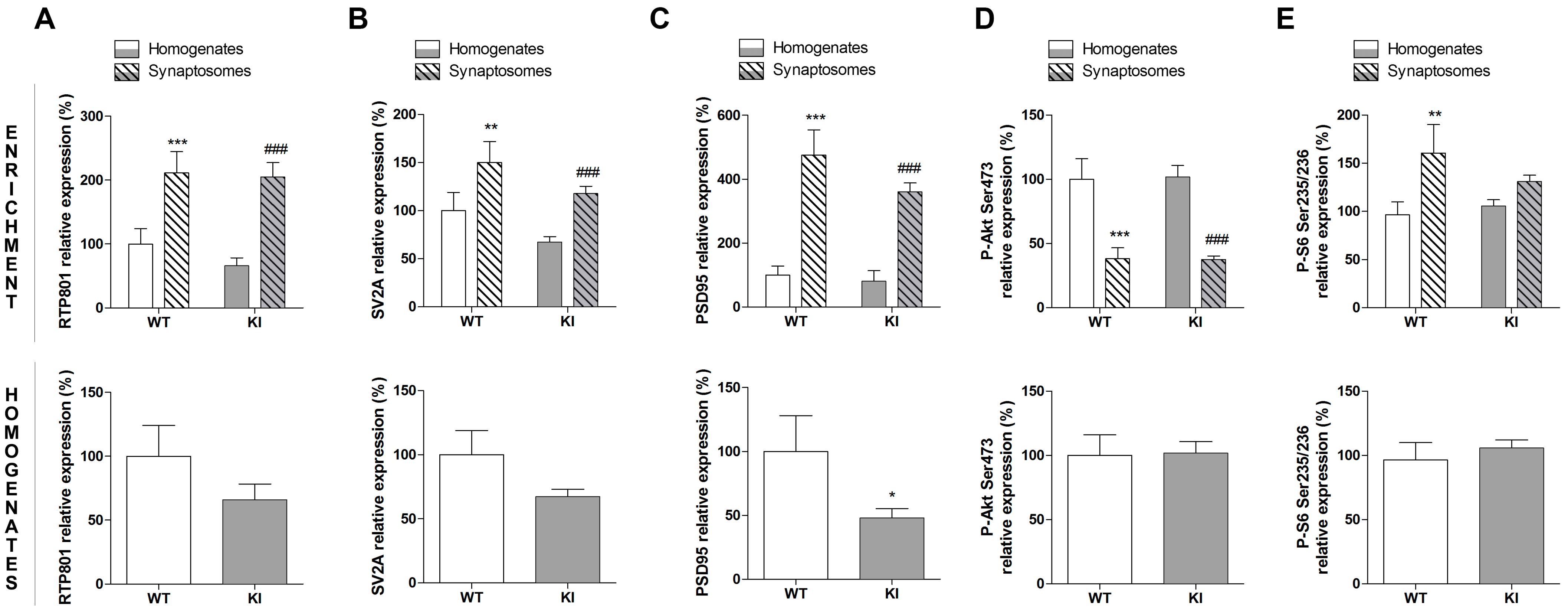

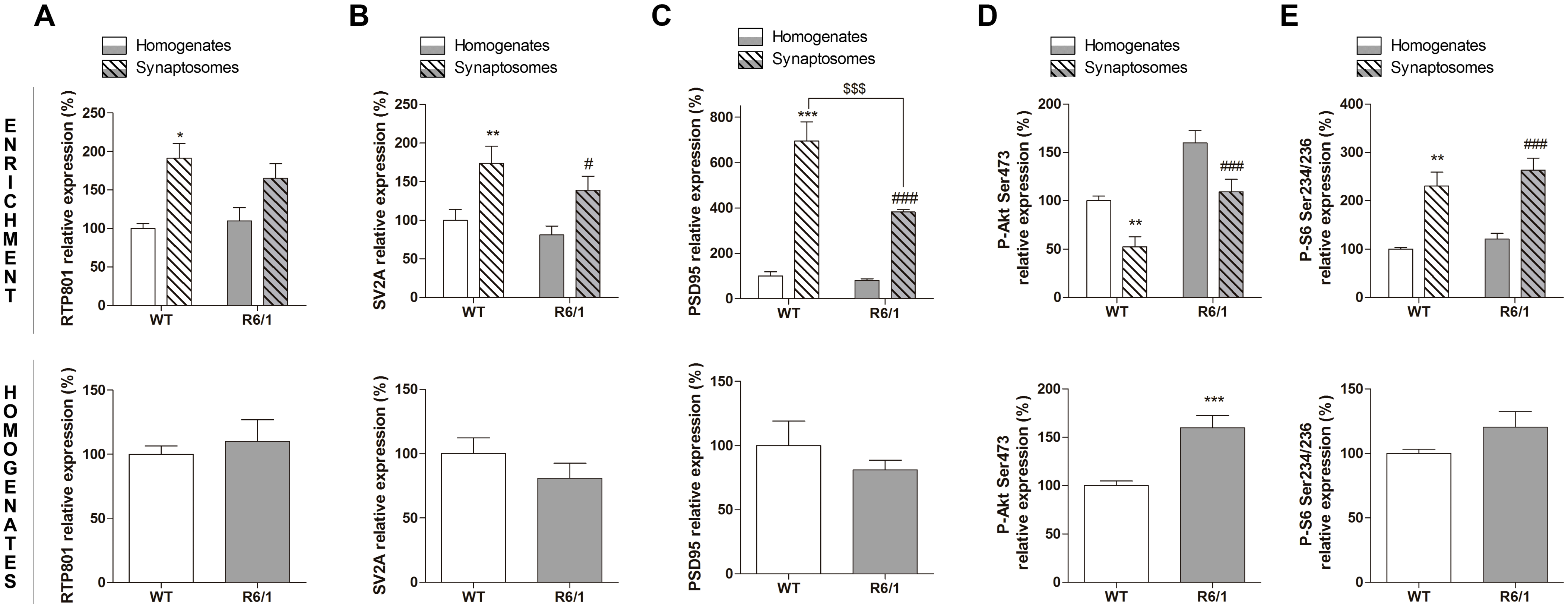

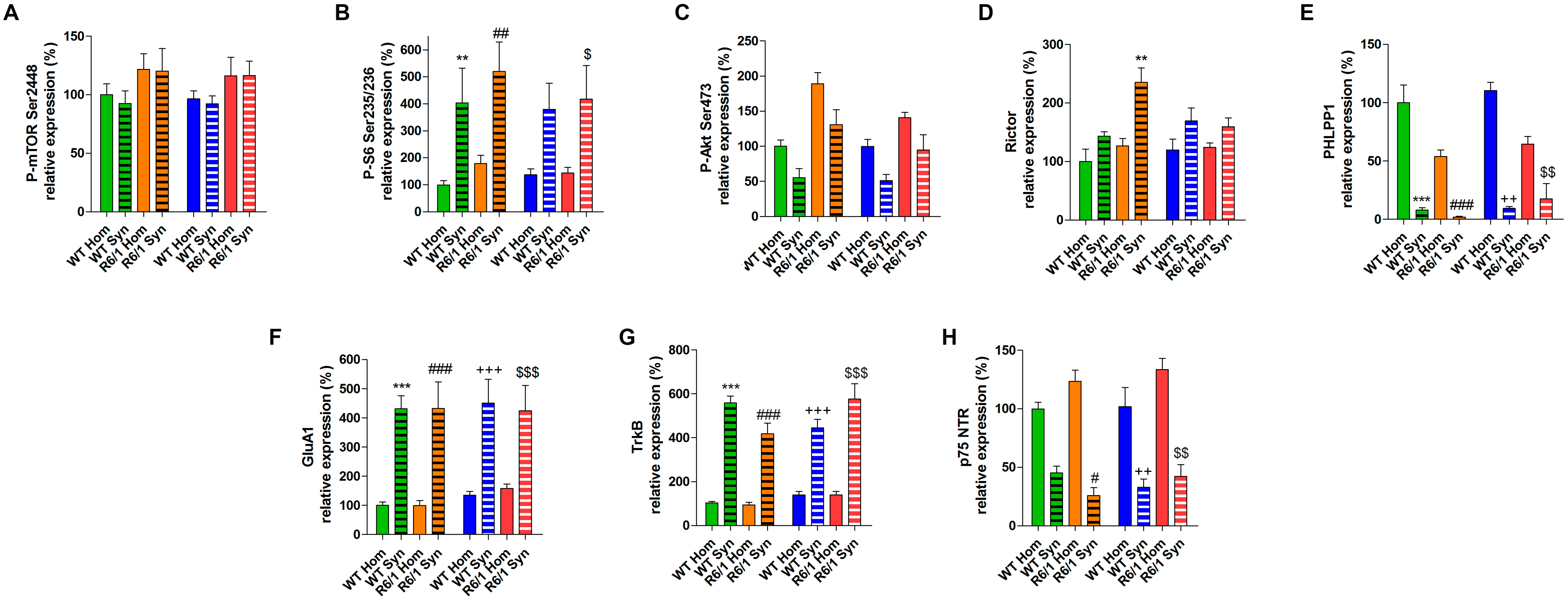
