## Supplementary table for "Synaptic RTP801 Contributes to Motor Learning Dysfunction in Huntington’s Disease"

| **Patient** | **Pathological diagnosis** | **Gender** | **Age (years)** | **CAG repetitions** | **Area analyzed** |
| --- | --- | --- | --- | --- | --- |
| 1 | Normal | Male | 83 | - | Str & Ctx |
| 2 | Normal | Male | 73 | - | Ctx |
| 3 | Normal | Male | 76 | - | Str & Ctx |
| 4 | Normal | Male | 64 | - | Str & Ctx |
| 5 | Normal | Male | 86 | - | Str & Ctx |
| 6 | Normal | Male | 58 | - | Str & Ctx |
| 7 | HD, Vonsattel grade 3-4 | Male | 55 | - | Str & Ctx |
| 8 | HD, Vonsattel grade 3 | Male | 53 | 45 | Str |
| 9 | HD, Vonsattel grade 1 | Male | 73 | 40 | Ctx |
| 10 | HD, Vonsattel grade 3 | Male | 85 | 40 | Str & Ctx |
| 11 | HD, Vonsattel grade 2 | Male | 76 | 41 | Str & Ctx |
| 12 | HD, Vonsattel grade 2 | Male | 72 | - | Str & Ctx |
| 13 | HD, Vonsattel grade 2-3 | Male | 68 | 42 | Str & Ctx |

**Supplementary Table 1.** Human post-mortem Huntington’s disease (HD) brains. Str = Striatum (Putamen); Ctx = Frontal Cortex.
